# Excitatory GABAergic modulation of calyx terminals in the vestibular sensory end organ

**DOI:** 10.1101/2020.04.30.069682

**Authors:** Yugandhar Ramakrishna, Soroush G. Sadeghi

## Abstract

GABAergic sources have been identified in the vestibular sensory neuroepithelium, mainly in the supporting cells. However, existence of GABA receptors or any possible GABAergic effects on vestibular nerve afferents has not been investigated. The current study was conducted to determine whether activation of GABA-B receptors affects calyx afferent terminals in the central region of the cristae of the semicircular canals in rats. We used patch clamp recording in P13 – P18 Sprague-Dawley rats of either sex. Application of GABA-B receptor agonist baclofen inhibited voltage activated outward potassium currents. This effect was blocked by selective GABA-B receptor antagonist CGP 35348. Antagonists of small (SK) and large (BK) current potassium channels resulted in an almost complete block of baclofen effect. The remaining baclofen effect was due to inhibition of voltage gated calcium channels and was blocked by cadmium chloride. Furthermore, baclofen had no effect in the absence of calcium in the extracellular fluid. Inhibition of potassium currents by GABA-B activation resulted in an excitatory effect on calyx terminal action potential firing. While in the control condition calyces could only fire a single action potential during step depolarizations, in the presence of baclofen they fired continuously during steps and a few even showed spontaneous discharge. We also found a decrease in threshold for action potential generation and a decrease in first spike latency during step depolarization. These results provide the first evidence for the presence of GABA-B receptors on calyx terminals, show that their activation results in an unusual excitatory effect and that GABA inputs could be used to modulate calyx response properties.

## INTRODUCTION

Vestibular end organs in the inner ear provide inputs to the vestibular nuclei in the brainstem through afferent vestibular nerve fibers. This information is used to provide vestibular reflexes that stabilize the gaze (the vestibulo-ocular reflex, VOR) and posture (the vestibulo-spinal reflexes) during daily activities. Particularly, afferents with more irregular resting discharge are shown to have high pass properties and carry information about fast head movements. The most irregular afferents receive inputs from type I hair cells through a specialized nerve terminal that covers the basolateral walls of the hair cell and are located in the central region of the cristae of the semicircular canals or maculae of otoliths (reviewed in: Goldberg 2000). In contrast, afferents with highly regular resting discharges receive inputs from type II hair cells through bouton terminals and are located in the most peripheral regions of the neuroepithelia. The majority of the afferents receive inputs from both type I and type II hair cells through calyx and bouton terminals, respectively. These ‘dimorphic’ afferents are located throughout the neuroepithelium. Furthermore, there is a direct relation between the number of bouton terminals and the regularity of the resting discharge of afferents (Goldberg et al. 1990; Holmes et al. 2017).

In recent years, the traditional idea that the vestibular periphery is a sensor with fixed properties has been challenged and it has been shown that the properties of hair cells and calyx terminals can be modified. Specifically, in all animals, an efferent pathway carries signals from the brainstem back to the end organs. In mammals, efferent neurons receive inputs mainly from the vestibular nuclei (Plotnik et al. 2002; Sadeghi et al. 2009). Efferent fibers project bilaterally to innervate bouton and calyx afferent terminals and type II hair cells. Interestingly, the effect of efferents on afferents is directly related to their resting discharge regularity, with the largest effects observed on the most irregular fibers (Goldberg and Fernandez 1980; Marlinski et al. 2004; Raghu et al. 2019; Sadeghi et al. 2009). Nicotinic (nAChR) and muscarinic (mAChR) acetylcholine receptors are present on type II hair cells and calyx afferent terminals and their activation results in changes in response properties of these cells (Holt et al. 2017; Holt et al. 2015; Kong et al. 2006; Kong et al. 2005; Lee et al. 2017; Li et al. 2007; Parks et al. 2017; Poppi et al. 2020; Poppi et al. 2018; Ramakrishna et al. 2020a; Zhou et al. 2020). In addition to changing the gain of the response (e.g., type II hair cell inhibition (Parks et al. 2017; Poppi et al. 2018; Zhou et al. 2020)), cholinergic inputs can inhibit potassium channels in afferents and affect their response properties (Perez et al. 2009; Ramakrishna et al. 2020a) and regularity of resting discharge (Kalluri et al. 2010).

The results of studies on gamma amino butyric acid (GABA) in the vestibular periphery are less conclusive. In mammals, GABA has been shown in efferent fibers (Kong et al. 1998a; b; Matsubara et al. 1995) and a recent study has found glutamic acid decarboxylase 67 (GAD67) – which is required for generation of GABA – and GABA itself, in supporting cells in mice vestibular end organs (Tavazzani et al. 2014). Another study in mice has found GAD2 in supporting cells and mainly hair cells in mice (Holman et al. 2019). While presence of GAD2 does not necessarily mean that GABA is produced, it indicates the possibility of having GABA in these cells. However, no previous study has investigated the possible effects of GABA on vestibular hair cells and afferents. In most brain areas and sensory organs GABA results in an inhibitory effect. Typically, activation of ionotropic GABA-A receptors result in activation of an inward chloride current and hyperpolarization of cells. Metabotropic GABA-B receptors are also known to provide inhibitory effects in most brain areas through pre and postsynaptic effects (reviewed in: Kantamneni 2016). A common presynaptic mechanism for the inhibitory effect of GABA-B receptor activation is inhibition of voltage gated calcium channels and a resultant decrease in vesicular release. Postsynaptic GABA-B receptors typically induce inhibitory effects through G-protein mediated pathways that activate GIRK and TREK-2 potassium channels, resulting in outward potassium currents and hyperpolarization of cells. Two exceptions have been reported where postsynaptic GABA-B activation results in an excitatory effect in the retina and substantia nigra. In the retina, this is mediated through a decrease in the activity of calcium channels, resulting in a decrease in BK potassium channel activity (Garaycochea and Slaughter 2016). In substantia nigra, GABA-B activation directly decreases the activity of SK potassium channels, through a cAMP/ PKA mediated pathway (Estep et al. 2016).

Previous studies have also shown the existence of different potassium channels that could be affected by GABA-B activation in the vestibular periphery, including inward rectifier K^+^ (Kir) (Udagawa et al. 2012), small current (SK) (Meredith et al. 2011), and large current channels (BK) (Limon et al. 2005; Schweizer et al. 2009). Here, we used whole-cell patch-clamp recording to investigate the presence of GABA-B receptors on vestibular afferent calyx terminals and whether they affected the activity of any of the potassium channels. We found that almost all calyces showed a GABA-B mediated response that consisted of a decrease in outward voltage sensitive potassium currents. GABA-B agonist (baclofen) application affected membrane properties to decrease spike frequency adaptation: calyx terminals that showed a single spike firing at the beginning of a step depolarization would fire many action potentials at a lower threshold in the presence of baclofen. Furthermore, the first spike latency decreased at the beginning of step depolarizations. We provide evidence that this effect is mediated through inhibition of voltage sensitive calcium channels and a resultant inhibition of BK and/or SK channels. These results provide the first evidence for the presence of GABA-B receptors on calyx terminals with an unusual excitatory effect and show that the response properties of calyx terminals can be modified by GABAergic inputs.

## MATERIALS AND METHODS

### Animals

Seventy two Sprague-Dawley rats (Charles River Laboratories), 13 to 18 days old and of either sex were used for the experiments. All animals were handled in accordance with animal protocols approved by the Institutional Animal Care and Use Committee at the University at Buffalo and carried out in accordance with NIH guidelines.

### Tissue preparation

Dissection of the end organ and tissue preparation were performed as previously described (Ramakrishna et al. 2020a; Sadeghi et al. 2014; Songer and Eatock 2013). Briefly, rats were decapitated after being deeply anesthetized by isoflurane inhalation. The skull was opened in the midline and the bony labyrinth was removed and placed in extracellular solution. Under the microscope, the ampullae of the horizontal and superior canals were opened. Attachments to bone were carefully dissected and the membranous labyrinth containing the cristae of the horizontal canal, anterior canal, and the macula of the utricle were taken out of the bone. The membranous labyrinth was opened over the two cristae and the utricle. The preparation was secured on a coverslip under a pin, transferred to the recording chamber, and perfused with extracellular solution at 1.5 −3 ml/min.

### Electrophysiology recording

The extracellular solution contained (in mM): 5.8 KCl, 144 NaCl, 0.9 MgCl2, 1.3 CaCl2, 0.7 NaH2PO4, 5.6 glucose, 10 HEPES, 300 mOsm, pH 7.4 (NaOH). The intracellular solution contained (in mM): 20 KCl, 110 K-methanesulfonate, 0.1 CaCl2, 5 EGTA, 5 HEPES, 5 Na2 phosphocreatine, 4 MgATP, 0.3 Tris-GTP, 290 mOsm, pH 7.2 (KOH). To perform patch-clamp recording, tissue was visualized with a 40x water-immersion objective, differential interference contrast (DIC) optics (Examiner D1 Zeiss microscope), and viewed on a monitor via a video camera (optiMOS, Qimaging). To make patch-clamp recording pipettes, 1 mm inner diameter borosilicate glass (WPI, 1B100F-4) was pulled with a multistep horizontal puller (P-1000, Sutter), coated with Sylgard (Dow Corning) and fire polished. Pipette resistances were 5-10 MΩ. All recordings were performed at room temperature (~ 23 °C). Application of drug solutions was performed (VC-6 channel valve controller, Warner Instruments, Hamden, CT) using a gravity-driven flow pipette (~100 μm tip diameter) placed at ~45o angle with the top of the crista near the area where a calyx was recorded from. The above solutions result in a liquid junction potential of −9 mV [50], which was corrected offline. Drugs were dissolved daily in the extracellular solution to their final concentrations from frozen stocks. R-baclofen (100 µM), CGP 35348 (300 µM), iberiotoxin (IBTX, 150 nM), apamin (Apa, 300 nM), cadmium chloride (CdCl, 0.5 – 1 mM), and XE-991 (10 µM) were purchased from Tocris Bioscience. Barium chloride (BaCl_2_, 100 µM) was purchased from Sigma.

### Experiment protocol

Whole cell patch clamp recordings were performed from calyx terminals in the central region of the cristae of anterior or horizontal canals. All measurements were acquired using Multiclamp 700B amplifier (Molecular Devices), digitized at 50 kHz with Digidata 1440A, and filtered at 10 kHz by pCLAMP10.7 software. Calyces innervated type I hair cells with typical morphology [50]. Cells were held at a holding potential of −70 mV resulting in a holding current of < 200 pA. To minimize time dependent changes (Hurley et al. 2006), all data were collected starting ~10 min after the membrane was ruptured. Since the effect of drugs increased with duration of application for the first ~10 min, to avoid time dependent changes in drug effects all data were collected 9-12 min after start of application (see: Ramakrishna et al. 2020a).

During voltage clamp recordings, a voltage step protocol was used to study calyx properties (Fig. 1A), which consisted of an initial holding potential of −79 mV (100 ms), a hyperpolarizing step to −129 mV (100 ms), 20 mV steps between −129 and +11 mV (300 ms), and return to holding potential of −79 mV. Depolarizing steps resulted in initial large Na+ inward currents and slower voltage sensitive outward (potassium) currents as previously described (Horwitz et al. 2014; Meredith and Rennie 2018; Ramakrishna et al. 2020a; Sadeghi et al. 2014; Songer and Eatock 2013). To study the effect of various drugs on responses to the voltage step protocol, current amplitudes were averaged for the final 100 ms of steps. Peak amplitude of tail currents were measured after returning to −79 mV at the end of steps and were plotted against holding potential at the preceding step (denoted as ‘activation step’ for x-axis title in figures).

**Figure 1.**
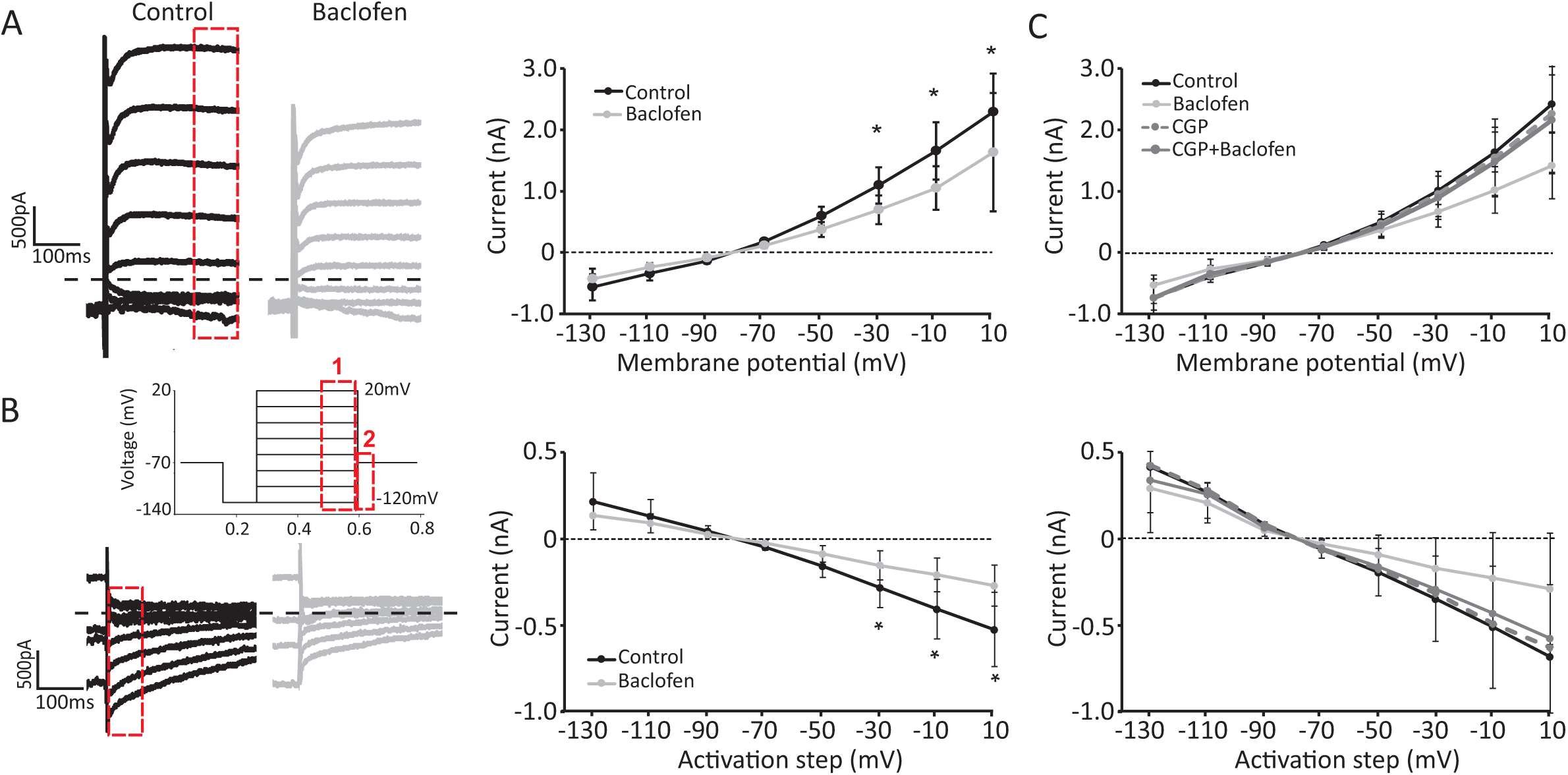
Activation of GABA-B receptors attenuates voltage‐activated currents. **(A)** Response of a calyx terminal to the voltage step protocol (20 mV steps between −129 mV and +11 mV) before (black) and after application of baclofen (grey). Calyx recording exhibited delayed rectifier type outward currents during depolarizing voltage steps (red box 1). Average responses show a significant decrease in voltage sensitive currents at the three most depolarized steps (marked by asterisk). Note, in all panels, changes in average currents are presented relative to the current at the initial holding potential of −79 mV. **(B)** The amplitude of tail currents after the steps decreased during baclofen application in an example recording (left, red box 2). Average tail currents (right) significantly decreased after the three largest depolarization steps (marked by asterisk). **(C)** Application of CGP 35348 (antagonist of GABA-B receptors) blocked the effect of baclofen, confirming that the effect is mediated through GABA-B receptors. All traces are from the same calyces recorded in the presence of different drug combinations.

During current clamp recordings, we first measured the resting membrane potential (i.e., 0 pA current injection). A current was then injected to hold the cells at −79 mV (for 5-10 s) and then current steps of 100 pA, 200 pA, 300 pA, 400 pA, and 500 pA (2 s duration) were injected with 3-5 s between different steps. None of the recorded calyces in the present study had spontaneous firing. The smallest of the five steps that generated an AP was considered as the AP generation threshold. The latency of the first spike was measured as the time from the beginning of the step to the peak of first AP for the threshold step. Finally, the number of APs during the steps were also counted. Results were compared before and after application of drugs.

### Data analysis

Clampfit 10 (Molecular Devices), Prism (Graphpad), and Matlab (Mathworks) softwares were used for analysis of data. Results are reported as mean ± SE. Paired t-test was used for comparison between two parameters and repeated measures two-way ANOVA with Bonferroni or Tukey post hoc test was used for comparisons of more than two conditions. Level of statistical significance was set at α = 0.05.

## RESULTS

Recordings were performed from the central zone of the crista of the horizontal or superior canals at postnatal days 13 – 18 so that type I hair cells and their calyx afferent terminals had acquired their characteristic morphological and electrophysiological properties (Favre and Sans 1979; Hurley et al. 2006; Lysakowski and Goldberg 2008; Rusch et al. 1998). Patch clamp recordings were performed from calyx terminals as previously described (Ramakrishna et al. 2020a; Sadeghi et al. 2014). The average resting membrane potential of calyces during whole-cell recordings was −71.9 ± 3.8 mV (n = 14) similar to that reported previously in mammals (Lim et al. 2014; Ramakrishna et al. 2020a; Sadeghi et al. 2014; Songer and Eatock 2013). Voltage step protocols were used to investigate voltage dependent changes in membrane currents (Fig. 1A). The effect of drugs were measured on currents at the last 100 ms of step depolarizations (Fig. 1A, red box 1) and on tail currents (Fig. 1A, red box 2). During depolarizing steps, calyces displayed outward currents mainly due to activation of delayed rectifier type currents by KCNQ and erg channels (Hurley et al. 2006; Lysakowski et al. 2011), calcium sensitive SK channels (Meredith et al. 2011), and probably voltage sensitive BK channels (Limon et al. 2005). After the depolarizing step, a tail current was observed most likely due to accumulation of potassium in part in the closed type I hair cell – calyx synaptic space (Contini et al. 2017; Lim et al. 2011).

### GABA-B activation suppressed voltage gated currents

To investigate whether GABA-B receptors are present on the calyx terminal and their possible effect on membrane properties of calyces, we applied R-baclofen, an agonist of GABA-B receptors during calyx recordings. Baclofen application (100 µM) resulted in a decrease in voltage dependent currents during depolarizing steps in every recorded calyx (n = 15), suggesting the presence of GABA-B receptors on these afferent terminals (Fig. 1A). For the population of recorded calyx terminals, application of baclofen resulted in an inhibition of currents for the more depolarized voltage steps > −29 mV (repeated measures ANOVA, p < 0.0001, Bonferroni posthoc test, p < 0.001 for −29 mV, −9 mV, and +11 mV). As expected, tail currents that are most likely due to the potassium accumulation (at least in part between the calyx and type I hair cell) during depolarization of the calyx were also suppressed during baclofen application (Fig. 1B). For the population of recordings, tail currents decreased following depolarizing steps of above −29 mV (repeated measures ANOVA, p = 0.0001, Bonferroni posthoc, p < 0.01 for −29 mV and p < 0.001 for −9 mV and +11 mV). The above changes were completely blocked in the presence of CGP 35348 (300 µM), a GABA-B antagonist. Figure 1C shows the average response for 4 calyx recordings in which voltage sensitive currents were inhibited with baclofen application. After washing baclofen, CGP 35348 was applied, which by itself had no effect on the currents in the same calyces. This was then followed by application of a combination of CGP 35348 with baclofen, which showed no effect (repeated measures ANOVA, p > 0.05 for CGP 35348 vs. CGP 35348 and baclofen conditions). These results strongly suggest that calyx terminals (at least in the central region of the cristae) have functional GABA-B receptors.

### Kir channels are not affected by GABA-B activation in calyx terminals

Previous studies have shown that GABA-B activation in different neurons and glial cells typically increases inward rectifying potassium (Kir) currents (O’Callaghan et al. 1996, Isomoto et al. 1997), including those mediated by Kir4.1 channels (Takeda et al. 2015). Kir4.1 channels are also expressed in calyx terminals (Udagawa et al. 2012) and it has also been proposed that similar to glial cells, they play a role in clearance of K^+^ from the closed synaptic space between type I hair cells and calyx terminals. Despite baclofen having a clear inhibitory effect on potassium currents at depolarized membrane potentials in the above experiments, we still tested whether GABA-B has any possible effect on Kir channel activity in the calyx. Voltage steps similar to those shown in Figure 1 were used before and after baclofen application in the absence and presence of barium chloride (BaCl_2_, 100 µM), a voltage independent blocker of Kir channels (O’Callaghan et al. 1996). We observed no change in baclofen effect on potassium currents in the presence of BaCl_2_ (n = 6, repeated measures ANOVA, p > 0.05), suggesting that Kir channels were not affected by GABA-B activation.

### GABA-B inhibits Ca^2+^-sensitive K^+^ channels through inhibition of voltage sensitive Ca^2+^ channels

Two recent studies have shown GABA-B mediated inhibition of BK (large current) potassium channels in the retinal ganglion cells (Garaycochea and Slaughter 2016) and SK (small current) channels in the substantia nigra (Estep et al. 2016). Since calyx terminals also express SK (Meredith et al. 2011) and probably BK (Limon et al. 2005) channels, we studied whether a similar GABA-B mediated inhibition was also present in the calyx terminal. We used a combination of antagonists for SK channel (apamin, 300 nM) and BK channel (iberiotoxin, 150 nM). As expected, the outward voltage dependent currents (Fig. 2A) and tail currents (not shown) were both inhibited in the presence of apamin and iberiotoxin. For the population of recorded calyces (n = 6) the outward current was decreased by 16% – 26% (for steps > −69 mV) with application of apamin and iberiotoxin. The decrease in voltage dependent currents were significant for most steps (Fig. 2B, repeated measures ANOVA, P < 0.006, post-hoc Tukey HSD, p > 0.8 for −129 to −69 mV and p < 0.03 for other steps). Adding baclofen resulted in a non-significant decrease in currents by ~10% (repeated measures ANOVA, p = 0.33). Similar effects were observed for tail currents at the end of voltage steps (not shown). To investigate whether the baclofen effect was due to inhibition of a voltage sensitive calcium current, we added XE-991 (KCNQ channel blocker) to the cocktail to inhibit all major voltage activated K^+^ currents in the calyx. Again, baclofen resulted in a small decrease in the average current (n = 7, largest decrease of 163 ± 115 pA for +11 mV step) (Fig. 2C). A similar decrease in currents was observed with addition of the nonspecific Ca^2+^ channel blocker cadmium chloride (CdCl_2_) (Fig. 2C), suggesting that baclofen inhibits voltage sensitive Ca^2+^ channels, similar to that shown in the retinal ganglion cells (Garaycochea and Slaughter 2016; Shen and Slaughter 1999).

**Figure 2.**
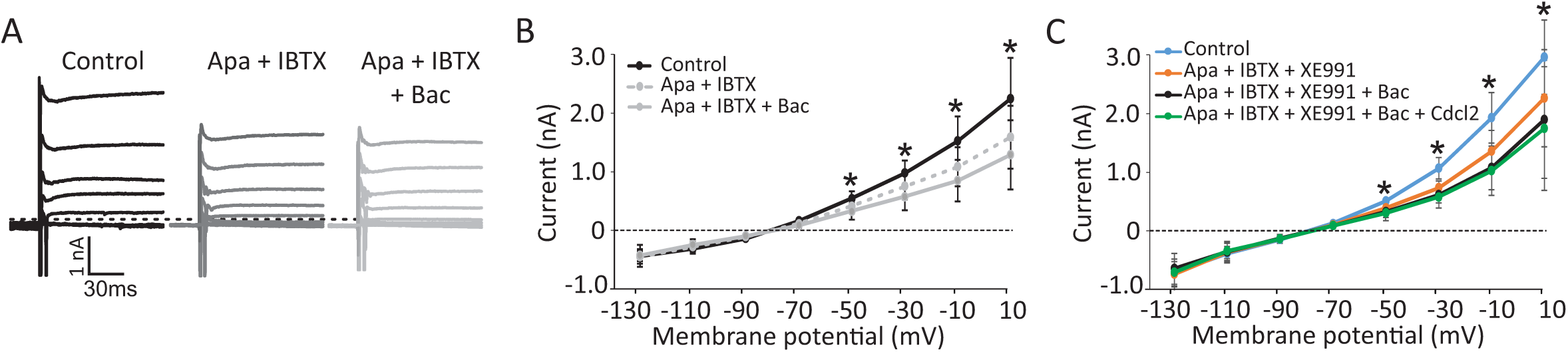
Effect of baclofen could be inhibited by blocking SK and BK channels. **(A)** Example calyx recording showing a decrease in outward currents during the step protocol in the presence of apamin (SK blocker) and iberiotoxin (BK blocker), which blocked the effect of baclofen. **(B)** Average of the recorded currents during the step protocol shows that apamin and iberiotoxin application blocked some of the voltage sensitive currents and that application of baclofen results in a small non-significant further decrease in the currents for the most depolarized steps. **(C)** Application of a cocktail of apamin, iberiotoxin, and XE-991 (KCNQ channel blocker) resulted in a decrease in voltage sensitive currents. Addition of baclofen resulted in a further small non-significant decrease in the currents at depolarized steps, which was blocked by cadmium chloride (a general blocker of calcium channels), suggesting that baclofen inhibits voltage sensitive calcium channels. * signify significant differences with control conditions (repeated measures ANOVA, posthoc Bonferroni test, p < 0.03).

We further explored the effect of GABA-B on Ca^2+^ channels in two sets of experiments. First, in the absence of calcium in the external solution (0 [Ca^2+^]), baclofen did not have any effect on the depolarizing steps and tail currents (Fig. 3A). Note that currents were decreased with 0 [Ca^2+^] due to the inhibition of voltage sensitive Ca^2+^ currents and Ca^2+^ activated potassium currents, but there was no further decrease with baclofen application. For the population of recordings in control external solution, 0 [Ca^2+^] external solution, and subsequent baclofen application (n = 7), voltage sensitive currents and tail currents decreased in 0 [Ca^2+^] compared to control, but there was no further change with baclofen application in 0 Ca^2+^ (Fig. 3B). In a second set of experiments in the control external solution, we applied CdCl_2_ to block calcium channels (n = 4), which decreased the amplitude of currents during depolarizing steps as well as the tail currents and occluded the effect of baclofen (Fig. 3C). In 3 recordings, we used CdCl_2_ with 0 [Ca^2+^] external solution to completely block all sources of external calcium to the cell. This resulted in inhibition of voltage activated and tail currents and complete block of the effect of baclofen (Fig. 3D). These results show a central role for Ca^2+^ in baclofen mediated effects and suggest that inhibition of voltage sensitive calcium channels by GABA-B receptors results in an inhibition of SK and/or BK channels.

**Figure 3.**
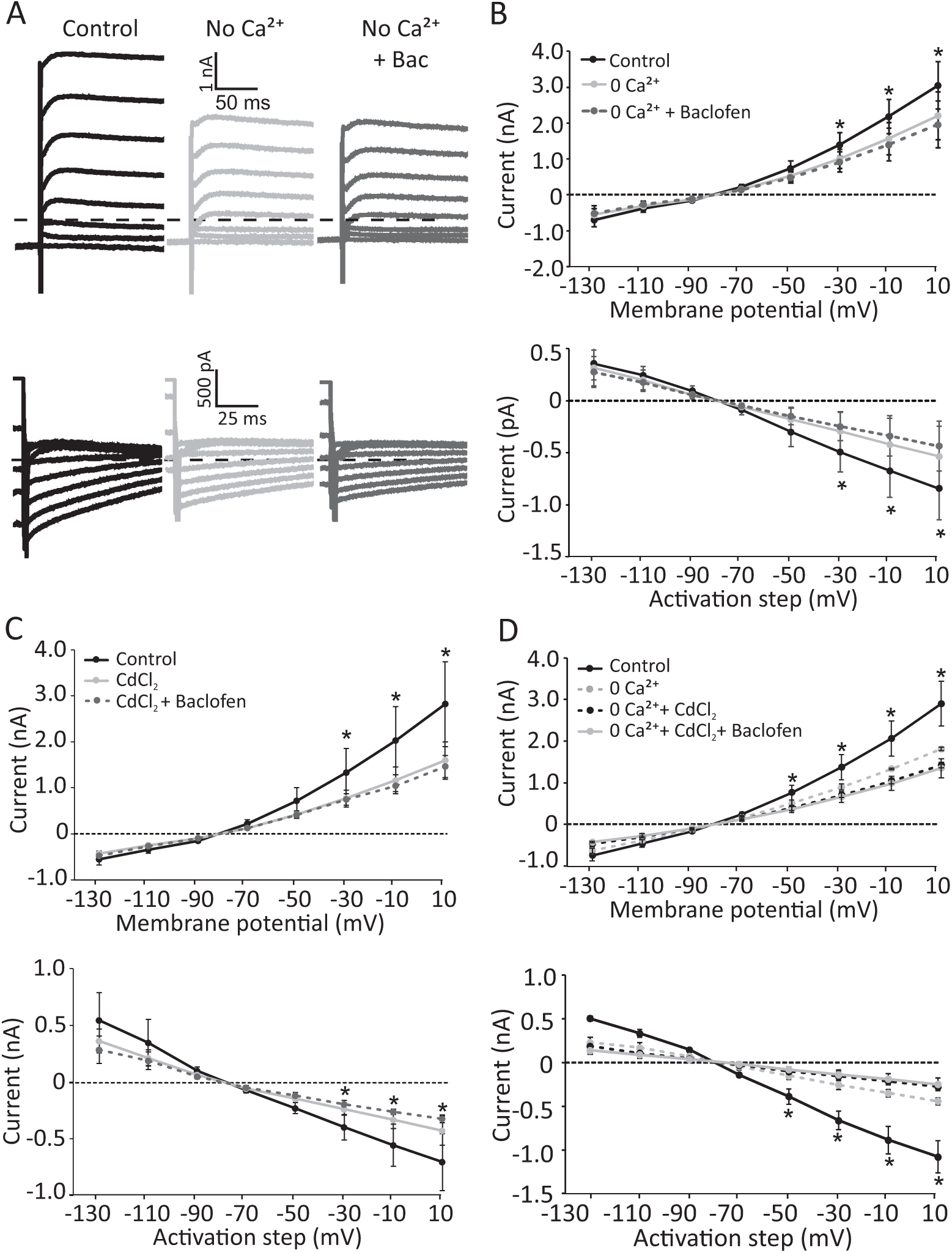
The effect of baclofen was mediated through calcium channels. **(A)** Example calyx recording showing a decrease in voltage sensitive currents during the step protocol and in tail currents in 0 [Ca^2+^] external solution, most likely due to inhibition of Ca^2+^-dependent K^+^ currents. The effect of baclofen was blocked under these conditions. **(B)** Average currents recorded from calyces during and after (tail currents) the step protocol. Application of 0 [Ca^2+^] external solution decreased the currents and almost completely blocked the effect of baclofen. **(C)** Application of calcium channel blocker cadmium chloride resulted in a decrease in voltage sensitive currents and tail currents and almost completely blocked the effect of baclofen, suggesting that baclofen functions through voltage sensitive calcium channels. **(D)** Application of the combination of 0 [Ca^2+^] external solution and cadmium chloride resulted in a complete block of baclofen effect on voltage sensitive currents and tail currents. * denotes significant differences with control condition (repeated measures ANOVA, posthoc Bonferroni test, p < 0.03).

### GABA-B activation has an excitatory effect on calyx terminals

We next used current clamp to investigate the effect of GABA-B activation on firing of action potentials (AP). For all recordings, we injected currents (if necessary) to have an initial calyx membrane potential at −79 mV. We then injected current steps of 100 – 500 pA in 100 pA steps. In control condition (i.e., before application of drugs), none of the recorded calyces (n = 15) showed spontaneous firing of APs and the threshold for generating an AP was reached by injecting 320 ± 147 pA. At threshold and more depolarized membrane potentials, all recorded calyces fired only a single AP at best, as previously reported (Ramakrishna et al. 2020a; Songer and Eatock 2013). The latency of this single spike was 6.8 ± 2.1 ms for the threshold current. After baclofen application calyces fired more than one spike during depolarization (Fig. 4A). On average, calyces generated more than 1 spikes after baclofen application compared to control condition for 200 pA, 300 pA, 400 pA, and 500 pA steps (Fig. 4B). The threshold for spike generation also significantly decreased (control: 320 ± 147 pA, baclofen: 219 ± 122 pA, paired t-test, p = 0.009) (Fig. 4C). Furthermore, the responses became faster with GABA-B activation as quantified by the first spike latency at the beginning of step depolarizations (Fig. 4D). For the threshold current, the latency showed a non-significant decrease (~2 ms) from 6.8 ± 2.1 ms to 5.0 ± 0.9 ms (paired t-test, P = 0.12). However, the latency of the first spike decreased with increasing amplitude of step currents (for the control condition as well as in the presence of baclofen) and the difference between the two conditions was significant for the 400 pA and 500 pA steps (paired t-test, p > 0.05 for 100 – 300 pA steps, p = 0.028 for 400 pA step and p = 0.003 for 500 pA step).

**Figure 4.**
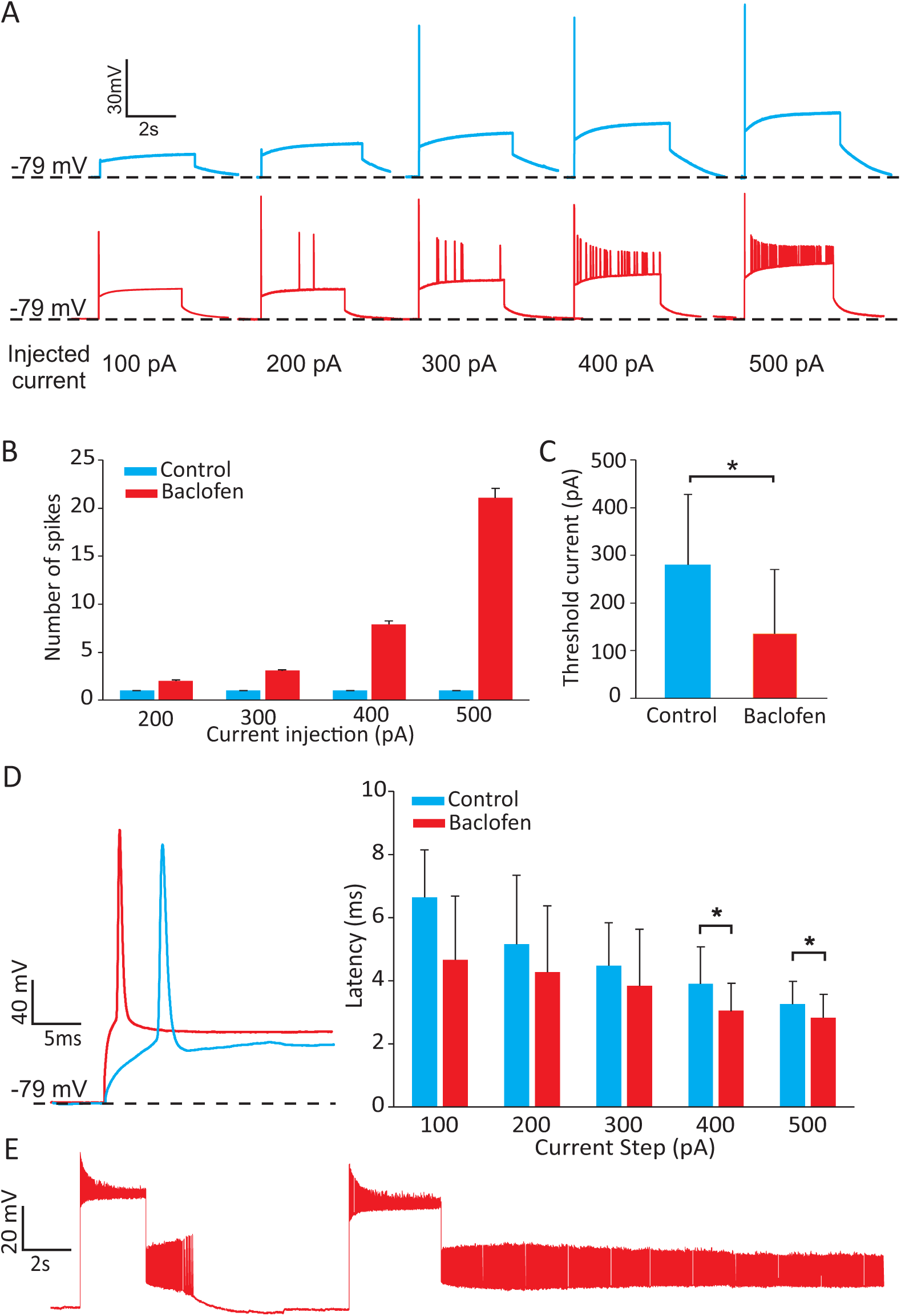
Calyces become more sensitive and respond faster following activation of GABA-B receptors. **(A)** Examples of current clamp recordings from calyces that at best fire a single AP in the control condition in response to injection of 100 – 500 pA current steps, but in the presence of baclofen fired many APs during the step. **(B)** The average number of spikes during steps were higher after baclofen application for all current steps. The control condition had 0 or 1 spikes for all steps, while after baclofen calyces fired at least 1 spike, even for the smallest current step. **(C)** The amplitude of the current required for AP generation (i.e., current threshold) decreased significantly during baclofen application. **(D)** The latency of the first spike decreased during application of baclofen compared to control condition. The decreases were significant for the two largest steps. **(E)** Some calyces showed spontaneous AP firing during baclofen application. In all panels control results are shown in blue and baclofen condition is in red; significant changes are marked by asterisks.

Previous studies have shown that both SK and BK channels affect neuronal spike frequency adaptation and action potential firing patterns in different brain areas and animal models (e.g., Deemyad et al. 2012; Deemyad et al. 2011; Gittis et al. 2010; Smith et al. 2002). Although we expected to see changes in firing patterns, it was surprising to see excitatory effects with GABA-B activation. Furthermore, while none of the recordings showed spontaneous AP firing in control condition, 8 out of 15 recorded calyces in current clamp exhibited spontaneous firing of AP in the presence of baclofen (Fig. 4E). Note that the other two reports of excitatory GABA-B effects in retinal ganglion cells and substantia nigra neurons did not show any spontaneous firing (Estep et al. 2016; Garaycochea and Slaughter 2016).

## DISCUSSION

In this study, we provide evidence for an unusual excitatory effect of GABA-B receptors on membrane properties of a group of sensory nerve terminals in the vestibular periphery. Application of GABA-B agonist baclofen resulted in inhibition of K^+^ currents at depolarized membrane potentials in calyx afferent nerve terminals. This effect was occluded by exhausting available Ca^2+^ sources through application of 0 [Ca^2+^] external solution and using calcium channel blocker CdCl_2_. Our results indicate that GABA-B acts through inhibition of voltage sensitive Ca^2+^ channels, resulting in inhibition of Ca^2+^ sensitive SK and/or BK potassium channels. This was further shown by a decrease in baclofen effect in the presence of SK and BK channel antagonists. These changes also resulted in faster responses (i.e., decreased first spike latency in response to step currents), increased sensitivity (i.e., decreased threshold), and increased gain (i.e., more spikes in response to depolarization). Notably, some of the calyx terminals showed spontaneous firing in the presence of baclofen, a property that is not typically seen in the *in vitro* preparation in the central regions of the cristae (Ramakrishna et al. 2020a; Sadeghi et al. 2014) and maculae (Songer and Eatock 2013). Note that this is independent of any synaptic inputs from hair cells to calyces, despite a decrease in K^+^ accumulation (as evidenced by decreased tail currents), and mainly due to changes in calyx membrane properties.

### Excitatory GABAergic activity in the vestibular periphery

Typically, activation of the metabotropic GABA-B receptors results in an inhibitory effect. On the presynaptic side, this is mediated by inhibition of voltage gated Ca^2+^ channels and a resultant decrease in synaptic vesicular release (reviewed in: Kantamneni 2016). On the postsynaptic side, inhibition is achieved through activation of K^+^ channels. Only two previous studies have reported an excitatory GABA-B mediated effect: (1) in retinal ganglion sensory neurons through inhibition of postsynaptic voltage sensitive Ca^2+^ channels resulting in inhibition of BK channels (Garaycochea and Slaughter 2016) and (2) in substantia nigra through inhibition of SK channels through a direct pathway involving adenylyl cyclase and protein kinase A (Estep et al. 2016). Our results indicate a mechanism similar to that described in the retina, however, because SK and probably BK channels are both present in the calyx, it is inevitable that both get affected by inhibition of voltage sensitive Ca^2+^ channels. This similarity suggests that such postsynaptic inhibition of Ca^2+^ channels might be a general mechanism for GABA-B action in peripheral sensory afferents and merit exploration in other sensory systems.

### Source of GABA in the vestibular periphery

Although previous studies have found GABA-A subunits on afferents in rats, mice, and hamsters (Foster et al. 1995; Kitahara et al. 1994) and our results provide ample evidence for the presence of GABA-B receptors on calyx terminals, a source of GABA in the vestibular periphery has not been unequivocally identified. There is evidence for existence of GABAergic efferent fibers and terminals in the peripheral vestibular neuroepithelium in mouse (Kong et al. 2002a), rats (Kong et al. 1998b; Matsubara et al. 1995), squirrel monkey (Usami et al. 1987), and humans (Kong et al. 1998a; Kong et al. 2002b). However, the efferent cell group in the brainstem cannot be labeled with anti-GABA antibodies (Perachio and Kevetter 1989). A recent study used GAD67-GFP or GAD65-GFP mice that express GFP along with the two isoforms of the enzyme glutamate decarboxylase (GAD), which transform glutamate to GABA (Tavazzani et al. 2014). This study concluded that only GAD67 is expressed in vestibular end organs and only in supporting cells in the peripheral regions of the semicircular canal cristae. No GFP labeled cells were observed in hair cells or in GAD65-GFP mice. Since GAD65 is believed to be responsible for generating GABA in synaptic vesicles (compared to GAD67 that generates cytoplasmic GABA) (Martin and Rimvall 1993; Soghomonian and Martin 1998) lack of this enzyme suggests that the generated GABA in supporting cells does not have vesicular release. Interestingly, using anti-GABA antibodies this study found that GABA was present in the supporting cells in both central and peripheral regions of the cristae. Another recent study used GAD2-tdTomato mice, which expressed red fluorescence with GAD2 and found fluorescence in both hair cells and supporting cells in the peripheral regions of the cristae and maculae (Holman et al. 2019). With increasing age, this expression was strongest in type I hair cells in the peripheral regions. This finding suggests presence of GABA as part of the afferent pathway, as has been suggested in frogs (Lopez and Meza 1988), guinea pig (Lopez et al. 1990; Meza et al. 1992), and toadfish (Holstein et al. 2004). However, presence of GAD2 does not necessarily indicate generation of GABA and it is unlikely that GABA is an afferent neurotransmitter AMPA receptor blockers suppress all synaptic events in calyx terminals (Sadeghi et al. 2014). From these previous studies, the most likely source for GABAergic inputs in the vestibular periphery are supporting cells. It is also possible that supporting cells provide non-vesicular release of GABA similar to glial cells (Attwell et al. 1993; Heja et al. 2009; Wu et al. 2007).

### Function of GABA-B activation in the vestibular periphery

Presence of different types of potassium (e.g., SK, BK, KCNQ, Kir, …) channels in calyx terminals decreases their membrane resistances, making it difficult to depolarize the calyx. We recently showed that muscarinic acetylcholine receptors (mAChR) inhibit KCNQ channels in calyx terminals, resulting in similar excitatory effects (Ramakrishna et al. 2020b), supporting the findings of previous studies in turtle (Holt et al. 2017). Inhibiting SK and BK channels have been shown to affect the spike frequency adaptation and firing patterns in other brain areas and animal models (e.g., Deemyad et al. 2012; Deemyad et al. 2011; Gittis et al. 2010; Smith et al. 2002). Interestingly, calyces in the central regions of the *in vitro* preparation of the vestibular neuroepithelium can only fire a single AP in response to a step depolarization. This could be due to the fact that efferent fibers are cut and separated from their cell bodies in the brainstem in this *in vitro* preparation and do not have spontaneous activity, as evidenced by the absence of any non-glutamatergic synaptic events in calyces (Sadeghi et al. 2014). This is further supported by our recent *in vivo* results that showed suppression of efferents results in a decrease or complete shutdown of mainly irregular afferents in mice (Raghu et al. 2019). Note that the lack of spontaneous firing can also be due to low release rates from hair cells onto the calyx, however, under the same conditions, changing the membrane properties of the calyx results in spontaneous firing of action potentials.

A second function for GABA-B receptors can be a gain control mechanism. Inhibition of SK/BK channels and the resulting increase in the number of APs during step depolarizations is akin to an increase in the response gain. Similar increases in the spontaneous firing and response gain have been observed in the vestibular nuclei following inhibition of BK channels (Nelson et al. 2017; Nelson et al. 2005; Smith et al. 2002). In addition to direct gain control, changes in the activity of potassium channels affect membrane filtering properties and its resonance at subthreshold potentials (Beraneck and Straka 2011).

### Interaction between cholinergic efferents and GABAergic cells

Finally, a recent study in the cochlea has shown expression of GABA-B receptors by efferent terminals that contact inner and outer hair cells (Wedemeyer et al. 2013). As a result, application of baclofen decreased the inhibitory cholinergic efferent inputs onto hair cells. Current evidence suggest that cholinergic efferents on calcyes are excitatory (Holt et al. 2017; Holt et al. 2015; Lee et al. 2017; Ramakrishna et al. 2020a) and any inhibition of efferent inputs should result in a decrease in calyx responses, which is the opposite of the effect observed here. However, it is possible that a similar effect is present in efferents that synapse onto type II hair cells, which are very similar to cochlear efferents in that they hyperpolarize type II hair cells through a9-containing receptors (Parks et al. 2017; Poppi et al. 2018; Zhou et al. 2020). Interestingly, in the vestibular periphery, supporting cells that contain GAD2 are activated by acetylcholine (Holman et al. 2019), which can play the role of a negative feedback control over cholinergic efferents.

### Conclusion

Overall, our results provide evidence for an unusual excitatory effect of GABA-B receptors on calyx terminals, which modulates their resting discharge, sensitivity, and membrane properties. This could potentially affect the response properties of irregular afferents, which play a major role in carrying information about fast head movements. These findings question the traditional notion that only central vestibular pathways are modulated during adaptation and compensation and suggest that GABAergic inputs along with cholinergic efferent inputs can provide the means for continuous fast adjustments of response properties of the vestibular periphery.

## ACKNOWLEDGEMENTS

This work was supported by NIDCD RO3 DC015091 grant and a Research Grant from the American Otological Society to SGS.

